# scCotag: Diagonal integration of single-cell multi-omics data via prior-informed co-optimal transport and regularized barycentric mapping

**DOI:** 10.64898/2025.12.11.693589

**Authors:** Yang Li, Yuting Tan, Rui Chen, Xin Maizie Zhou, You Chen, Yuankai Huo, Xingyi Guo, Anshul Tiwari, Zhexing Wen, Xue Zhong, Bradley A. Malin, Bingshan Li, Zhijun Yin

**Affiliations:** Department of Computer Science, Vanderbilt University, Nashville, Tennessee; Department of Molecular Physiology and Biophysics, Vanderbilt University, Nashville, Tennessee; Department of Biomedical Engineering, Vanderbilt University, Nashville, Tennessee; Department of Biomedical Informatics, Vanderbilt University Medical Center, Nashville, Tennessee; Department of Electrical and Computer Engineering, Vanderbilt University, Nashville, Tennessee; Department of Pathology, Microbiology, and Immunology, Vanderbilt University Medical Center, Nashville, Tennessee; Division of Epidemiology, Department of Medicine, Vanderbilt Epidemiology Center, Vanderbilt-Ingram Cancer Center, Vanderbilt University Medical Center, Nashville, Tennessee; Department of Psychiatry and Behavioral Sciences, Emory University, Atlanta, Georgia; Department of Medicine, Vanderbilt University Medical Center, Nashville, Tennessee; Department of Biostatistics, Vanderbilt University Medical Center, Nashville, Tennessee

**Keywords:** Diagonal integration, Multiomics, Optimal transport, Representation learning

## Abstract

Recent advances in high-throughput single-cell technologies have enabled characterization of cellular states across distinct omics layers, yielding complementary insights into the organization of biological systems. To jointly leverage these disparate modalities, computational methods have been developed to align non-overlapping cell populations profiled in different omics layers, a task known as diagonal integration, thereby facilitating more fine-grained biological interpretations. However, existing integration approaches unrealistically assume that all cells are alignable and treat prior-knowledge-derived feature correspondences as fully reliable, retaining them unrefined throughout the alignment process. In this work, we introduce *scCotag*, a co-optimal transport (COOT)-based deep learning framework for diagonal integration of single-cell multi-omics data, to address these limitations. *scCotag* first infers cell alignment and feature correspondence with prior-informed COOT in an iterative manner, and then leverages the resulting transport plans to jointly learn cell and feature embeddings via regularized barycentric mapping and graph reweighting. Systematic benchmarking on human brain, bone marrow, and blood single-cell RNA-seq and ATAC-seq datasets demonstrates the overall superior performance of *scCotag* over state-of-the-art methods in both cell alignment accuracy and embedding accuracy. In particular, *scCotag* excels in simulated imbalance scenarios where not all cells are alignable. Applying *scCotag* to data in postmortem brains of Alzheimer’s disease (AD) and non-AD (NoAD) patients further yields more fine-grained biological interpretation of regulatory mechanisms. Together, these results demonstrate that *scCotag* provides a robust framework for single-cell diagonal integration and regulatory inference.

## 1 Introduction

Advancements in high-throughput single-cell technologies have revolutionized our understanding of cellular mechanisms. These technologies, including transcriptomics, proteomics, metabolomics, epigenomics, etc., describe complementary facets of cell states, thus together enabling finer interpretations than any individual modality alone [20]. However, due to the destructive nature of these technologies [6], it may not always be possible to profile two or more modalities simultaneously in the same cell population. Although cutting-edge sequencing protocols have now enabled the joint profiling of multi-omics data [7,34], their application remains constrained by high costs and technical complexity [3]. To exploit the vast amount of existing unpaired multiomics data for more fine-grained biological interpretation (e.g., gene regulatory network (GRN) inference [5]), it is crucial to computationally align cells representing the same underlying biological state across different omic layers. This problem, known as diagonal integration, presents a unique challenge since no anchored cells or shared features are available [2].

Several computational methods for diagonal integration have been developed [17]. The predominant strategy relies on biological prior knowledge to link features across omic layers, thereby imputing one modality (e.g., gene-activity scores) from the other (e.g., chromatin-accessibility peaks) [24,36]. However, priors derived from external paired datasets [15] or from genome coordinates [19,37] are inherently context-dependent and noisy, thereby compromising alignment accuracy. Alternatively, some methods directly infer cell correspondences from the intra-modality geometric structure, under the assumption that a shared latent manifold structure is preserved between modalities [25,14,12]. In practice, this assumption is frequently violated because real-world datasets often contain modality-unique cell types and imbalanced compositions [28,41]. Deep learning-based methods extend these ideas by learning low-dimensional cell representations and projecting them into a modality-shared latent space for cell alignment [8,9,33,38,23]. These cell representations support various downstream analyses, including but not limited to clustering [33], cross-modality imputation [11], and label transfer [9]. Nevertheless, these approaches still suffer from noisy priors and restrictive assumption that all cells are alignable, providing limited mechanisms to prevent over-alignment of modality-specific populations.

Here, we introduce *scCotag*, a deep generative framework that integrates unpaired **s**ingle-**c**ell multi-omics data via prior-informed **co**-optimal **t**ransport (COOT) and regularized b**a**rycentric mappin**g** to address these limitations. scCotag explicitly incorporates dynamic biological priors into COOT [39] to obtain transport plans, i.e., probabilistic measures quantifying the cross-modality correspondences at both cell- and feature-levels, that respect both prior knowledge and intra-modality geometry. Leveraging these transport plans, scCotag introduces a novel transportation-guided representation learning framework that (i) denoises a biological prior graph through graph reweighting using the feature-level transport plan, and (ii) aligns cells across modalities via barycentric mapping using the cell-level transport plan. To handle realistic imbalance scenarios where not all cells are alignable, scCotag further exploits within- and cross-modality geometric structure to estimate alignment confidence and uses anchor-based regularization to align only the reliably alignable cells by enforcing the geometric preservation. Systematic benchmarking on three paired multi-omics datasets from distinct organs demonstrates that the scCotag overall outperforms state-of-the-art methods in both alignment accuracy and embedding accuracy. In simulated imbalance settings, scCotag maintains high alignment accuracy on alignable cells while minimizing geometric distortion for unalignable ones. Finally, applying scCotag to unpaired multi-omics data from postmortem brains of Alzheimer’s disease (AD) and non-AD patients yields biologically meaningful and disease-relevant gene–peak regulatory interactions, illustrating its utility for regulatory inference from unpaired single-cell datasets.

## 2 Methods

### 2.1 Overview of scCotag

scCotag is a two-stage deep generative framework for diagonal integration of single-cell multi-omics data (Fig. 1). The first stage aims to obtain biologically meaningful cross-modality correspondences at both cell and feature levels. To this end, scCotag extends COOT [39] by directly encoding biological prior knowledge into the OT kernel and iteratively updating these priors with the data-refined transport plans, namely prior-informed COOT. It yields cell- and feature-level transport plans that respect both known biological priors and the intrinsic geometry within each modality.

**Fig. 1.**
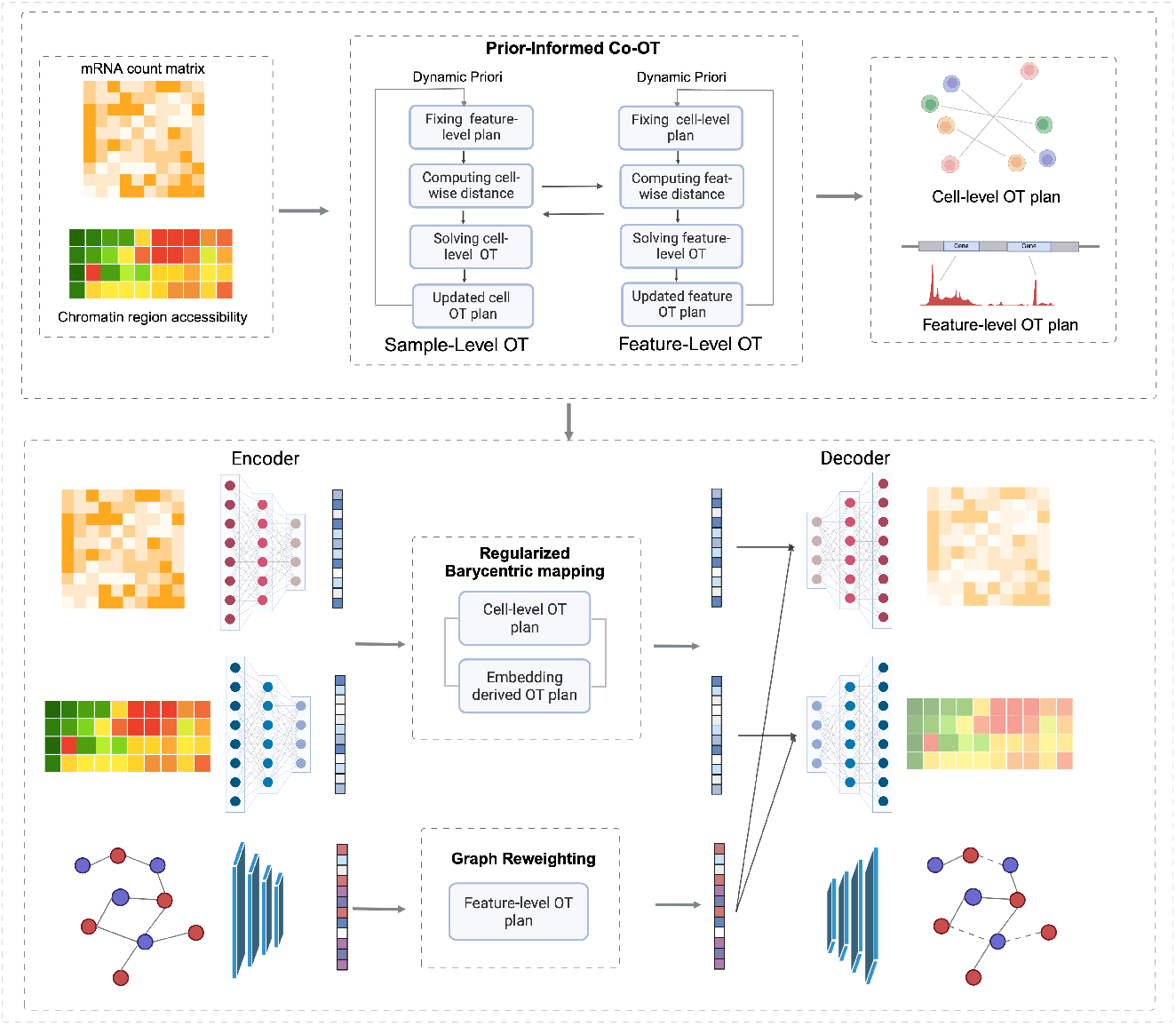
Overview of the two-stage scCotag framework. **Top:** A prior-informed COOT iteratively infers cell-level and feature-level transport plans from two unpaired modalities and updates the priors with plans obtained in the previous iteration. **Bottom:** Modality-specific VAEs learn cell embeddings from the input omic layer through regularized barycentric mapping. The VGAE learns feature embeddings on a biological prior graph with graph reweighting. The learned embeddings are combined to reconstruct the original omics profiles. *Figure created with* https://BioRender.com.

Although the cell transport plan achieves high alignment accuracy, it cannot be utilized for downstream tasks that require cell representations. Thus, the second stage of scCotag jointly learns cell and feature representations by leveraging the obtained transport plans. Specifically, at the cell level, scCotag projects cells into a modality-shared latent space via regularized barycentric mapping with the cell-level transport plan. At the feature level, it denoises biological prior graph via graph reweighting using the feature-level transport plan. In this way, scCotag learns biologically meaningful cell and feature representations that can be directly applied to various applications.

A major challenge in diagonal integration is the imbalance across modalities, marked by unbalanced batch size, unbalanced cell type proportions, and modality-unique cell types [29,42], which contaminate the downstream analysis. Existing approaches [33,8] leverage unbalanced optimal transport (UOT) [32,10] to tackle the imbalance scenarios where not all cells are alignable, which relaxes the mass constraint, allowing for the detection of outlier cells by partially destroying mass in the transport plan. In practice, however, we observed that identifying unalignable cells (outliers) is far more challenging than identifying high-confidence alignable cells. Motivated by this observation, scCotag estimates an *alignment confidence* for each cell from within- and cross-modality geometric structure. High-confidence anchor cells are identified and used to regularize the barycentric mapping, encouraging low-confidence cells to preserve their relative geometric structure with respect to all anchors. This design mitigates geometric distortions under imbalance while maintaining high alignment accuracy on alignable populations.

Together, these components collectively enable scCotag to achieve robust and geometry-preserving alignment across different omic layers (see **Appendix C** for the ablation study). More details regarding model design are provided below.

### 2.2 Prior-informed co-optimal transport

Given two modalities, 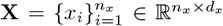 and 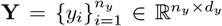 the classic OT aims to find a transport plan *P* that moves one modality onto the other at minimum cost [35,6]. Formally,

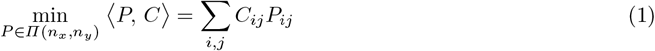

where *C* is the cost function measuring the pairwise distance between cells in **X** and **Y**, and Π(*n*_*x*_, *n*_*y*_) is the set of probability measures that enforce the balanced transport plan, i.e., 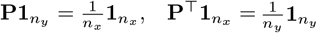. Nevertheless, solving equation 1 is computationally expensive and often yields sparse transport plans. Entropic-regularized OT (EOT) [13] extends the classic OT by adding an entropy term weighted by *ε* (see equation 2), offering smoothness, computational efficiency, and numerical stability, thus being favored in real-world applications, where *H*(*P* ) =∑ _*i,j*_ −*P*_*ij*_ (log *P*_*ij*_ − 1 ).

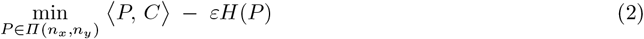

However, EOT assumes that the two point clouds lie in a shared feature space, which is invalid in diagonal integration. Gromov-Wasserstein OT (GWOT) [30] addresses cross-modality matching by penalizing intra-modality geometry distortion but fails to exploit the feature correspondence. COOT [39] overcomes this limitation by jointly inferring the correspondences between samples and features (equation 3), where ℒ measures the pairwise distance, *P*_*s*_ is the sample-level plan, and *P*_*v*_ is the feature-level plan.

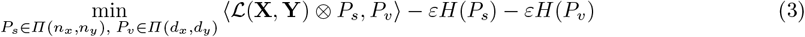

Nonetheless, vanilla COOT does not incorporate biological prior knowledge, which is crucial for multi-omics data integration [9,37,36]. In scCotag, we explicitly encode such priors into the COOT kernel *K* by replacing the smoothness-guiding uniform distribution with a prior *Q* as follows:

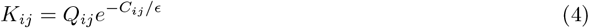

which encourages the obtained transport plan to stay close to *Q* while still adapting to the data. We construct different priors for cells and features. The cell-level initial prior 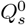 captures cross-modality cell correspondence derived via EOT on the RNA count matrix and the imputed gene activity matrix. The feature-level prior 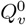 is a binary matrix encoding overlaps between extended gene bodies and chromatin accessibility peaks. In addition, scCotag iteratively updates the prior *Q* with the obtained transport plans from the previous iteration rather than using a fixed prior, allowing the priors to become increasingly data-refined. In summary, we formulate a prior-informed COOT as follows, where *t* indicates the *t-th* iteration:

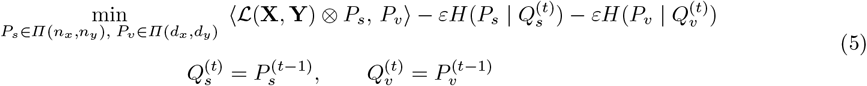

### 2.3 Learning cell and feature representations with transport plans

#### Graph reweighting

In scCotag, we utilize two modality-specific variational autoencoders (VAEs) [21] to extract cell representations, retaining the heterogeneous information within each modality. However, separately trained VAEs project cells into distinct latent spaces, making them incomparable. To link them, we follow the strategy proposed in [9] by introducing a prior graph 𝒢= (𝒱, ℰ ), where 𝒱 denotes the feature set and ℰ is the edge set. The prior graph is constructed from the feature pairs in feature-level prior 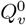 and used for feature representation learning through a variational graph autoencoder (VGAE) [22]. Since the prior graph 𝒢 contains noise (i.e., not all edges are real gene-peak interactions), we leverage the feature plan *P*_*v*_ to de-noise 𝒢 via a weighted graph reconstruction, which down-weights the importance of feature pairs receiving lower OT mass in *P*_*v*_. The importance of an arbitrary edge *w*_*i,j*_ is defined as follows:

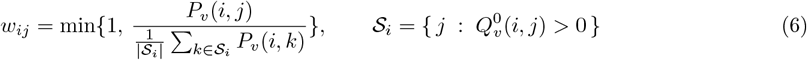

We then incorporate this weighted graph reconstruction into the evidence lower bounds (ELBO) of the VGAE as the reconstruction term, formulated in equation 7, where **v** represents the learned feature representations, *A* is the adjacency matrix derived from the prior graph 𝒢, *q*_*ϕ,graph*_(**v** | 𝒢) denotes the graph encoder parameterized by *ϕ, p*_*θ,graph*_(*A*_*ij*_ | **v**) represents the graph decoder parameterized by *θ*, and the sets ℰ_+_ and ℰ_−_ correspond to the positive and sampled negative edges, respectively.

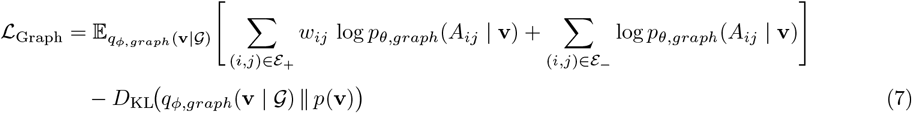

The learned feature representations **v** are applied to omics VAEs to reconstruct the omic profiles. We thus formulate the ELBO for omic VAEs in equation 8, where *q*_*ϕ,omics*_(**u**|**x**) denotes the encoder parameterized by *ϕ, p*_*θ,omics*_(**x**|**u, v**) represents the decoder parameterized by *θ*, and **u** denotes the learned cell embeddings:

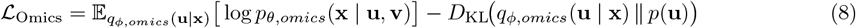

#### Regularized barycentric mapping

Given the cell-level transport plan *P*_*s*_ obtained in the first stage, we utilize it to refine the cell embeddings via barycentric mapping. To this end, we combine *P*_*s*_ with an embedding-derived transport plan *P*, which is computed from the current cell embeddings using a backpropagation-enabled entropic OT solver [16], weighted by a hyperparameter *λ*_*prior*_:

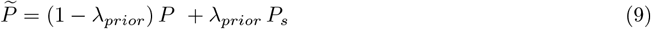

This mixed transport plan 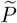 is subsequently used to perform barycentric mapping. During this mapping, we use one modality as the anchor and update only the other. This asymmetric updating strategy prevents training instabilities that may arise when both modalities are simultaneously updated. Let **X** be the anchor modality and **Y** be the updating modality. The barycentric mapping is formulated in equation 10, where **u**_*x,i*_ denotes the cell representation for *i* ∈ **X** and **ū**_*y,j*_ represents the barycentric image for *j* ∈ **Y**.

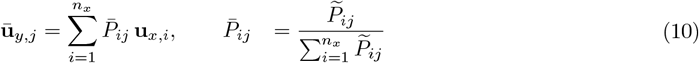

Each barycentric image is a weighted average of anchor embeddings determined by the normalized mixed transport plan 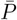. Finally, we encourage the cell embeddings of updating modality **Y** to align towards their barycentric projections and in turns, the anchor modality **X**, through a mean squared error loss:

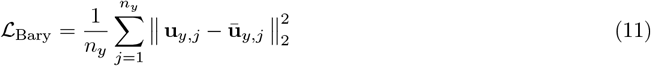

In imbalanced settings where not all cells are alignable, the vanilla barycentric mapping may indiscriminately align all cells despite their alignability. To mitigate this, we extend the barycentric mapping in equation 11 with a regularization term. Unlike traditional K-Nearest Neighbors (KNN)-based approaches that preserve local geometry within limited neighborhoods, this regularization term enforces the preservation of intra-modality global structure beyond local neighborhoods, thereby leaving modality-specific cells unaligned (see ablation results in **Appendix C**). Concretely, scCotag estimates an *alignment confidence* **c** ∈ (0, 1) for each cell in **Y** based on both within-modality and cross-modality geometry. Let *D*_within_ denote the within-modality distance matrix and *D*_cross_ the cross-modality distance matrix. The cross-modality proximity *s*_cross_ measures how close cells are, on average, to their nearest neighbors in the other modality, while the within-modality entropy *s*_within_ quantifies structural uncertainty within **Y** via a normalized entropy:

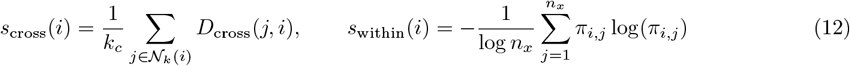

where *π*_*i,j*_ denotes the normalized similarity between cells *i, j* ∈ **Y** derived from *D*_within_, and 𝒩_*k*_(*y*) represents the set of the top *k*_*c*_ nearest neighbors of cell *i* across modalities. We then fuse *s*_within_ and *s*_cross_ and apply a robust z-score normalization with median absolute deviation, followed by a sigmoid transformation to yield the *alignment confidence* **c**. Formally,

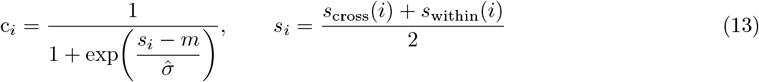

where *m* denotes the median value of *s*_*i*_, and 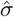 denotes the median of the deviations of *s*_*i*_. To distinguish alignable cells from uncertain ones, we model the distribution of **c** using a two-component Gaussian mixture model (GMM). Using the Bayes decision point as the threshold, cells with a confidence score above this threshold are recognized as high-confidence anchor cells (𝒜), while the remaining cells are classified as low-confidence non-anchor cells (𝒩). Given these two sets, we regularize the barycentric mapping by enforcing the relative distances between each non-anchor cell and all anchors remain consistent before and after projection. Let *D*_*u*_ denote the pairwise distances between cell embeddings **u**_*y*_, and *D*_*ū*_ the corresponding distances between their barycentric projections **ū**_*y*_. The anchor-based regularization loss is

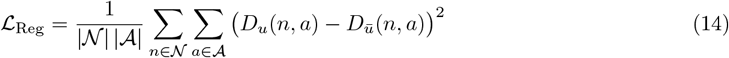

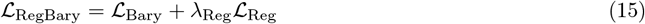

Finally, the overall training objective of scCotag is formulated as below, where ℒ_*Div*_ is the sinkhorn divergence encouraging distributional alignment between modalities **X** and **Y**:

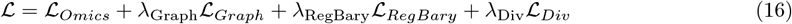

## 3 Experiment Settings

### 3.1 Datasets

To evaluate model performance, we used three real-world paired multi-omics datasets: peripheral blood mononuclear cells (**PBMC**) dataset [1], bone marrow mononuclear cells (**BMMC**) dataset [27], and a human brain dataset with Alzheimer’s Disease cells (**Tsai**) [26]. All three datasets provide joint profiling of gene expression and chromatin accessibility from the same cells, offering a ground truth for cell-level correspondence across modalities.

The PBMC dataset comprises 9,625 cells after standard preprocessing and quality control. The BMMC dataset contains 4,000 cells randomly sampled from the original dataset. The Tsai dataset includes multi-regional brain cells from participants in the Religious Orders Study/Memory and Aging Project (ROSMAP); we randomly sampled 10,000 cells and retained 5,604 cells after quality control. For PBMC and Tsai, gene activity matrices were imputed from fragment files using ArchR [19], whereas the BMMC dataset provides preprocessed gene activity scores.

To systematically study the effects of imbalance, we simulated three imbalance scenarios from the PBMC dataset with various degrees of imbalance. We first downsampled 100 cells per cell type to exclude the cell type proportion imbalance, resulting in a perfectly paired simulation dataset with 1,600 cells across 16 cell types in each modality. We then randomly removed 2, 4, and 6 non-overlapping cell types from each modality to generate datasets with increasing degrees of modality-unique cell types, denoted by *Simu1, Simu2*, and *Simu3*, respectively.

### 3.2 Evaluation metrics

We assessed model performance on three paired datasets in terms of two criteria, *cell alignment accuracy* and *embedding accuracy*, following the benchmarking paper [17]. The alignment accuracy was evaluated at both the single-cell and cell-type levels using *Fraction of samples closer than the true match* (**FOSCTTM**) and *cell-type match* (**CTM**). **FOSCTTM** quantifies the average proportion of cells that are closer than their true counterparts in the other modality. A score of 0 indicates perfect alignment. **CTM** measures the proportion of cells whose nearest neighbors across modalities share the same cell type; a higher CTM score indicates more accurate cell-type alignment.

To evaluate how well the learned cell embeddings remove batch effects while preserving biological structure [17], we used *adjusted mutual information* (**AMI**), *batch average silhouette width* (**batch-ASW**), and *graph connectivity* (**GC**). **AMI** measures the agreement between clusters derived from the learned cell embeddings and ground-truth cell type annotations. A higher AMI score indicates better preservation of biological identity. **Batch-ASW** measures how well cells from different modalities are mixed in the shared latent space while a higher score indicates better modality mixing. **GC** quantifies the average percentage of connected cells from the KNN graph in all cell types; a higher GC score indicates better connectivity and modality mixing.

When modality-unique cell types exist, an ideal integration method should accurately align only *alignable* cells, while leaving the modality-unique cell types unaligned. Current evaluations, however, are limited to alignable cells only and largely ignore the unalignable cells [33,8]; such unalignable cells, if erroneously aligned, can be detrimental for downstream analysis. To address this gap, we introduce two metrics tailored to imbalanced settings: *within-modality distortion* (**WD**) and *cross-modality distortion* (**CD**). **WD** measures the extent to which the intra-modality geometric structure of modality-unique cells is distorted after integration, by comparing distance patterns under imbalanced and perfectly paired settings. Formally,

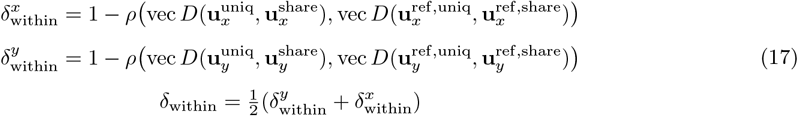

where *D* denotes the pairwise Euclidean distance, vec( · ) flattens a matrix into a vector, and *ρ* is the Pearson correlation. Superscripts *uniq* and *share* indicate modality-unique and shared subsets, while the superscript ref denotes the perfectly paired setting. **CD** is defined similarly by replacing within-modality distances with cross-modality distances, thereby quantifying distortion of cross-modality geometry.

### 3.3 Training details

We trained scCotag in two stages, consisting of a warm-up stage followed by a tuning stage. For preprocessing, we performed dimension reduction by applying PCA on scRNA-seq data and LSI on scATAC-seq data. The resulting 100-dimensional representations were used as the input to modality-specific VAEs. Each omics VAE encoder contains a two-layer neural network (256-256) with LeakyReLU activation. The decoder reconstructs the raw count matrix via a dot product between cell embeddings and feature embeddings, using a negative binomial distribution. The VGAE encoder initializes feature representations based on node indices and updates them using a graph convolutional network, while the decoder uses a dot product and models edges with a Bernoulli distribution. Since cell embeddings of different modalities lie in different feature spaces at the early training stage, we optimized only omics VAEs and the VGAE at the warm-up stage, 1000 epochs by default. In the following tuning stage, we introduced the regularized barycentric mapping loss and sinkhorn divergence loss with dynamically increasing weights according to the epoch, for a total of 3000 epochs. In both stages, we utilized the Adam optimizer with a learning rate of 1e-3 and a weight decay of 1*e* − 4. The default hyperparameters are as follows: *ϵ* = 0.05, *λ*_*Graph*_ = 1, *λ*_*RegBary*_ = 0.1, *λ*_*Div*_ = 0.01 and *λ*_*Reg*_ = 0.1.

### 3.4 Downstream task

One major application of integrating transcriptomic and epigenomic profiles is to decipher gene-peak cisregulatory interactions and infer regulatory networks. Therefore, we applied scCotag to an unpaired single-nucleus RNA-seq and ATAC-seq profiling dataset [40], including both AD patients and normal controls.

We grouped cells based on their pathology state (i.e., AD or NoAD) and sampled 3,000 cells via stratified random sampling by major cell types, curating one group with AD onset, denoted by the *AD* set, and the other group without AD, denoted by the *NoAD* set.

After training scCotag separately on the *AD* and *NoAD* sets, we computed pairwise gene-peak cosine similarity with the learned feature embeddings and conducted permutation tests with Benjamini-Hochberg correction [4] at a significance level of 0.05, to identify significant interactions. To assess the biological plausibility of inferred interactions, we used the gene-peak pairs reported in [40] as a reference for enrichment analysis.

## 4 Results

### 4.1 scCotag outperforms the state-of-the-art methods in systematic benchmarking

We benchmarked scCotag against five high-impact diagonal integration methods, including GLUE (*v0*.*4*.*0* ) [9], scConfluence (*v0*.*1*.*1* ) [33], uniPort (*v1*.*3* ) [8], MOSCOT (*v0*.*4*.*3* ) [23], and LIGER iNMF (*v2*.*2*.*1* ) [24], using their recommended preprocessing pipelines and default hyperparameters. Among these methods, GLUE serves as the current state-of-the-art diagonal integration method [17]. Fig. 2 summarizes the benchmarking results across all datasets and metrics (see **Appendix A** for details).

**Fig. 2.**
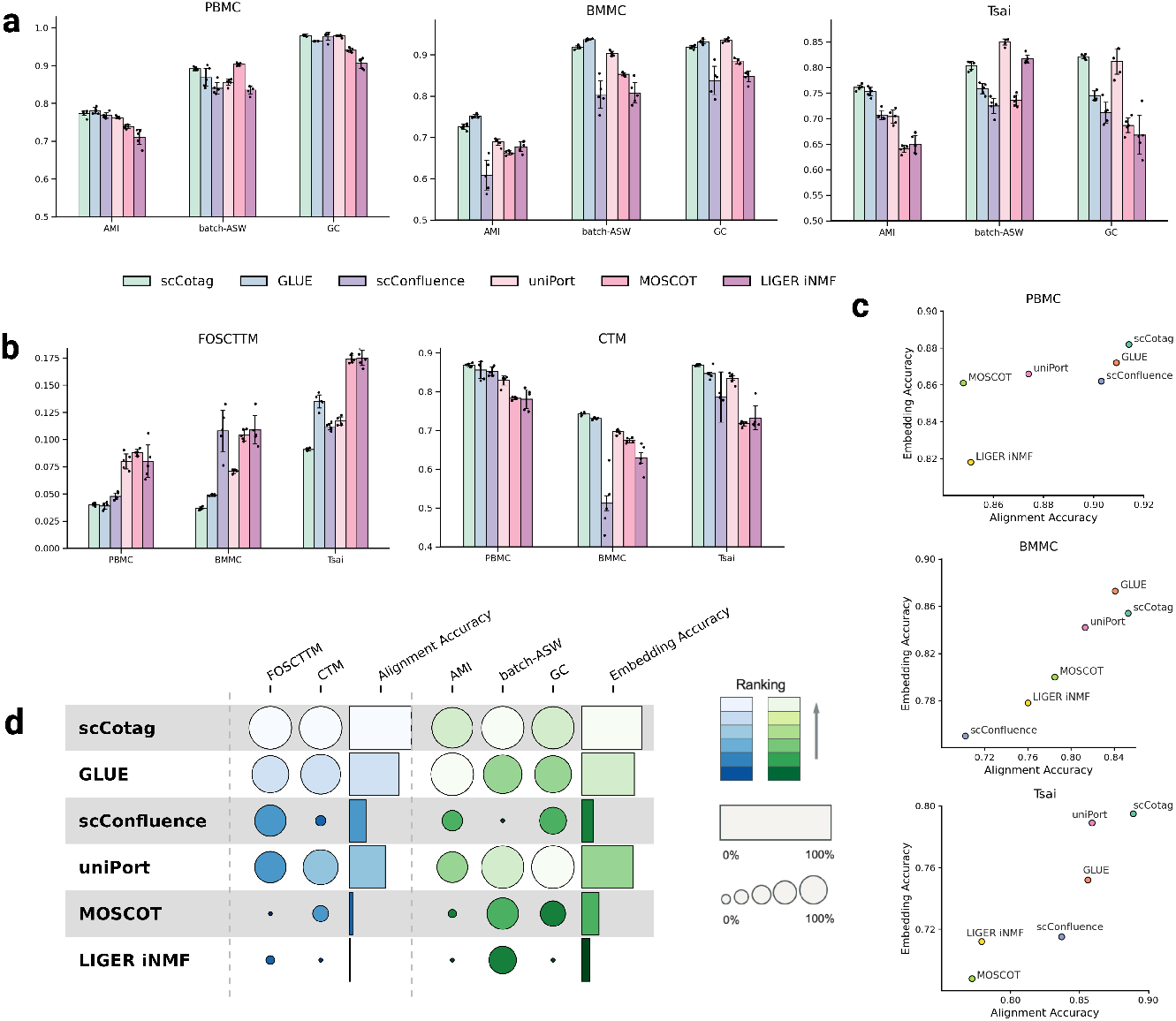
Benchmarking results on paired datasets. (**a**) Embedding accuracy evaluated by AMI, batch-ASW, and GC across three datasets (all: higher is better); dots denote results from individual random seed and error bars indicate standard deviation across seeds; (**b**) Alignment accuracy assessed by FOSCTTM (lower is better) and CTM (higher is better) across three datasets; (**c**) Comparison of embedding accuracy (defined as *Mean(AMI + batch-ASW + GC)*) and alignment accuracy (defined as *Mean((1-FOSCTTM) + CTM)*) on each dataset; (**d**) Overall performance, aggregated across datasets and within each metric using min-max normalization. The circle size reflects relative magnitude, and color intensity indicates performance ranking. We replicated the experiment five times with different run seeds and reported the mean values for each method.

Across all datasets, scCotag consistently achieves the highest or near-highest score on all three embedding accuracy metrics, demonstrating its robustness in preserving biological structure and removing modality effects (Fig. 2a). The aggregated results of embedding accuracy show that scCotag occupies the upper-right corner of the performance space on the PBMC and Tsai datasets, and exhibits competitive performance with GLUE on the BMMC dataset (Fig. 2c). As for the cross-modality alignment accuracy, scCotag consistently outperformed all competing methods in alignment accuracy, demonstrating the lowest (or equally lowest) FOSCTTM and the highest CTM scores across all three datasets (Fig. 2b,c). Notably, scCotag exhibited significant alignment accuracy improvements over existing methods in the Tsai dataset, which is characterized by lower read depth per cell, underscoring the robustness of scCotag in real-world integration scenarios where data quality is compromised.

We further assessed the overall performance of each method by averaging model performance across the three datasets and applying min-max normalization within each metric, following [42] (Fig. 2d). scCotag scored the highest on FOSCTTM, CTM and batch-ASW as well as near-highest in AMI and GC. Moreover, we averaged the normalized FOSCTTM and CTM scores into an alignment accuracy score, and normalized AMI, batch-ASW and GC into an embedding accuracy score. scCotag ranks first in both criteria, followed by GLUE and uniPort. In summary, these results demonstrate that scCotag achieves overall superior performance to current state-of-the-art methods in terms of both alignment accuracy and embedding accuracy.

### 4.2 scCotag preserves both within- and cross-modality geometry on imbalance scenarios

To evaluate model performance under imbalance scenarios where modality-specific cell types exist, we examined alignment accuracy for *alignable* cells (FOSCTTM) and geometry preservation for *unalignable* cells (within- and cross-modality distortion). Fig. 3 presents the model performance on the three metrics across three simulated imbalance datasets (see **Appendix B** for details). As imbalance becomes more extreme, existing methods exhibit a clear trade-off: methods such as GLUE and LIGER iNMF keep relatively low FOSCTTM but suffer from rapidly increasing distortion, whereas scConfluence and uniPort mitigate geometric distortion at the expense of alignment accuracy. In contrast, scCotag maintains the lowest (or near-lowest on *Simu1* ) FOSCTTM and the lowest within- and cross-modality distortion across all three simulations, indicating that it aligns shared populations without compromising the geometric structure.

**Fig. 3.**
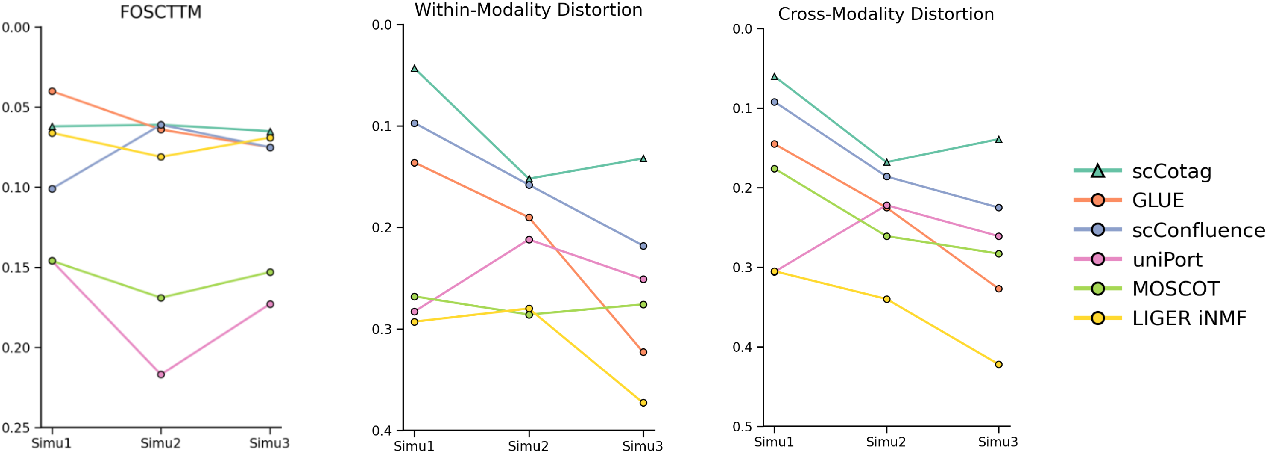
Model performance comparison under simulated imbalance scenarios. Model performance evaluated by FOSCTTM, within-modality distortion, and cross-modality distortion across three simulated imbalance scenarios, including *Simu1, Simu2*, and *Simu3*. We repeated the experiment five times with different run seeds and reported the median values given the high variability across different seeds under imbalance scenarios.

The UMAP visualizations in Fig. 4 confirm these behaviors in the *Simu2* setting. GLUE, MOSCOT and LIGER iNMF frequently over-integrate modality-unique populations, aligning them into clusters of different cell types (red circles in column 2), which explains their high distortion. scConfluence and uniPort suffer less from this over-alignment through UOT but leave substantial separation even among shared cell types (red circles in column 3), consistent with their reduced alignment accuracy. By contrast, scCotag yields well-aligned shared cell types with good cross-modality mixing, while keeping modality-specific populations distinct. Together, these results demonstrate that scCotag maintains high-accuracy alignment for alignable cells while effectively leaving modality-unique cells unaligned through its anchor-based regularization.

**Fig. 4.**
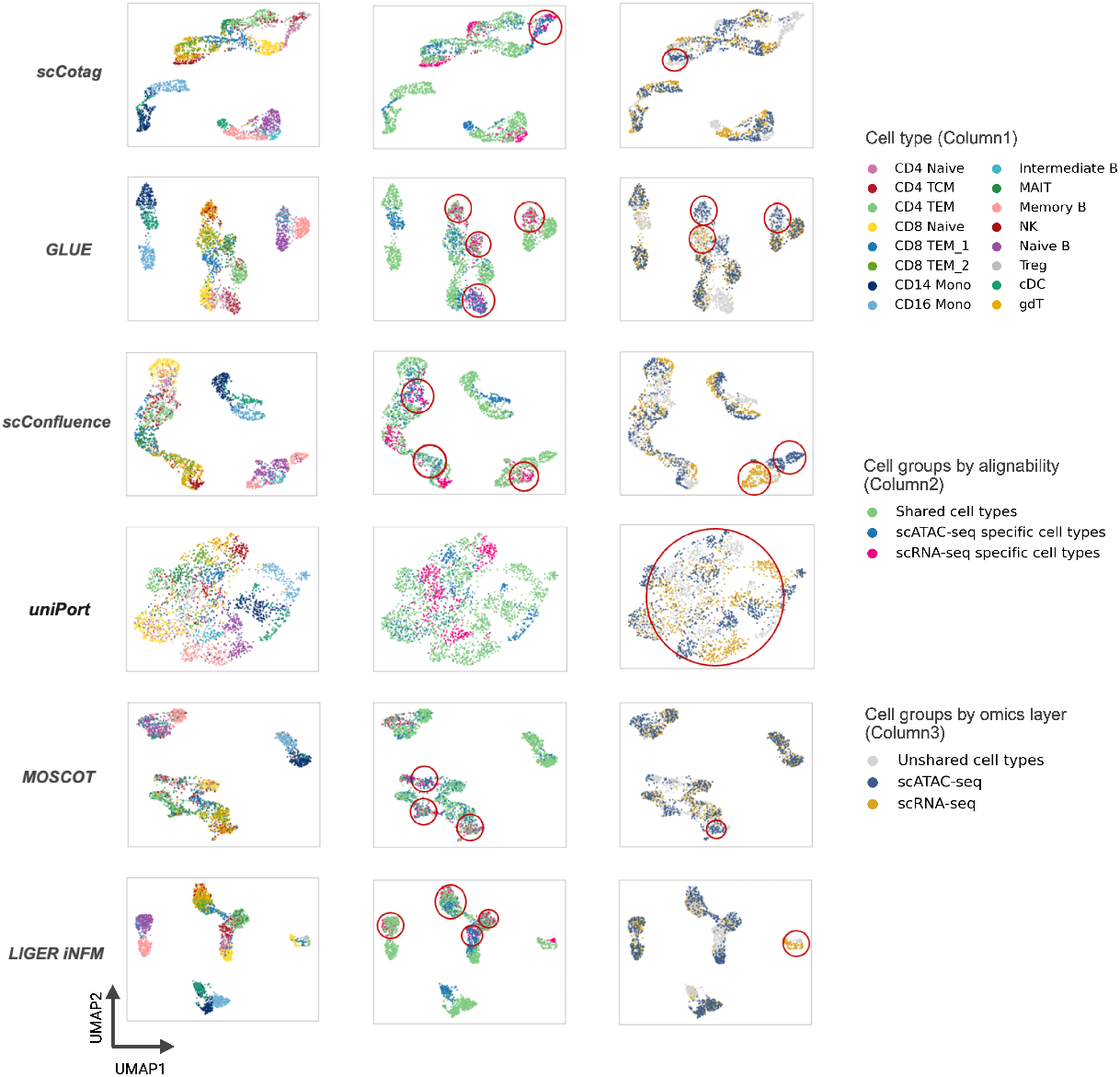
UMAP visualization of different model results on Simu2. Ideally, there should be no overlap between cell groups in the second column, and there should be perfectly mixed modalities in the third column. Incorrect overlaps in the second column or separations in the third column are highlighted with red circles.

### 4.3 scCotag performs regulatory inference on unpaired Alzheimer’s Disease data

We then applied *scCotag* for regulatory inference on the *AD* set and *NoAD* set, using the peak–gene scoring and significance procedure described in Section 3.4. We next tested whether significant peak–gene pairs were enriched for an external reference set of regulatory interactions. The enrichment test indicated that the significant pairs inferred by scCotag were substantially more enriched for the reference pairs than the non-significant ones (*p <* 1*e*^−300^, Fisher’s exact test), demonstrating scCotag’s ability in performing reliable regulatory inference using unpaired data.

Fig. 5 highlights a notable example. *THSD7B* has been identified as a down-regulated gene in vulnerable neurons in AD [18]. In the NoAD group, scCotag identifies regulatory interactions between *THSD7B* and nearby regulatory elements, whereas some of these interactions are absent in the AD group. Further mapping demonstrated that many of these NoAD-specific links (e.g., *THSD7B* -chr2:136665698-136666198, chr2:136874242-136874742 and chr2:137339039-137339539) overlap with distal enhancer-like signature candidate cis-regulatory elements (cCREs-dELS) [31], consistent with its down-regulation in AD given enhancer loss. These results also align with recent findings of epigenomic erosion in late-stage AD [26]. Moreover, several inferred interactions intersect fine-mapped eQTL regions (*r*^2^ *>* 0.6), providing additional support for the inferred regulatory interactions (Fig. 5). Together, these results demonstrate that scCotag not only integrates multi-omic profiles but also prioritizes disease-relevant regulatory interactions from unpaired single-cell datasets.

**Fig. 5.**
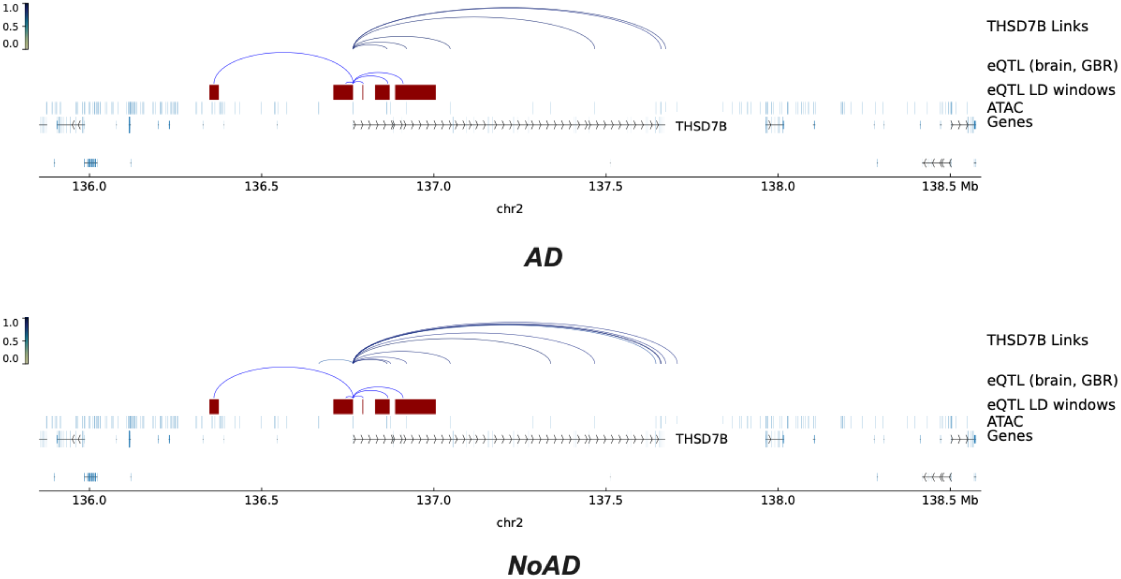
Inferred regulatory interactions between THSD7B and its regulatory elements by scCotag in the AD set (**top**) and NoAD set (**bottom**). Fine-mapped eQTL regions are highlighted in red.

## 5 Discussion and Conclusion

In this paper, we introduced scCotag, a two-stage deep generative framework for diagonal integration of single-cell multi-omics data. scCotag extends COOT with dynamic, biologically informed priors to jointly infer cell- and feature-level transport plans, and then uses these plans to guide representation learning through graph reweighting and regularized barycentric mapping. This design allows scCotag to leverage both modality-specific heterogeneity and cross-modality correspondence, producing cell and feature embeddings that are directly useful for downstream analyses.

Systematic benchmarking across multiple datasets demonstrated that scCotag achieves overall superior alignment and embedding quality compared with current state-of-the-art methods. Importantly, to better characterize integration quality under imbalance scenarios, we introduced imbalance-aware metrics that quantify within- and cross-modality geometric distortion. Under simulated imbalance scenarios with modality-unique cell types, scCotag maintains high alignment accuracy for alignable cells and the lowest geometric distortion for unalignable ones, whereas existing methods tend to either over-align unalignable cells or sacrifice alignment accuracy. These results highlight the effectiveness of our alignment confidence estimation and anchor-based geometry regularization in aligning only reliably matchable cells. Beyond integration, we applied scCotag to unpaired Alzheimer’s disease single-cell datasets, yielding biologically interpretable regulatory interactions pertaining to AD pathogenesis that are also consistent with enhancer erosion and fine-mapped eQTLs. Taken together, our results show that scCotag provides a robust and flexible framework for diagonal integration of single-cell multi-omics data. By jointly refining biological priors and representations while explicitly accounting for imbalance, scCotag brings multi-omic integration closer to the complexity of real-world datasets and enables reliable reconstruction of disease-relevant regulatory landscapes.

Despite promising results, we acknowledge several limitations that can serve as a basis for future work. First, the current implementation does not yet scale to atlas-level integration due to the computational cost of OT. Future extensions will incorporate mini-batch OT to improve its scalability. Second, our experiments focused only on scRNA-seq and scATAC-seq integration, but the framework is flexible and could be easily extended to other modalities (e.g., methylation or spatial transcriptomics) by adapting the prior construction. Third, scCotag explicitly targets one of the key imbalance scenarios of modality-unique cell types. Future work may evaluate how other types of imbalance affect alignment quality and downstream analyses.

## Code Availability

The source code is available at https://github.com/YangLi122/scCotag.

## Acknowledgments

This work was partially supported by the National Institute on Aging of the National Institutes of Health under award numbers R01AG069900, R01AG089717, and R01AG065611.

## Appendix A: model performance on paired datasets

We assessed model performance of scCotag and competing methods across three real-world paired datasets: PBMC (Table 1), BMMC (Table 2), and Tsai (Table 3). Specifically, we also reported the results for COOT and prior-informed COOT to validate the effectiveness of incorporating priors into OT kernels. Since the cell transportation plan cannot be utilized to compute embedding accuracy, we leave them as *Not applicable*, denoted by *NA*. We replicated the experiment five times with different random seeds and reported the mean and standard deviation as mean ± standard deviation. We evaluate alignment accuracy using FOSCTTM (lower is better) and CTM (higher is better), while embedding accuracy is measured via AMI, batch-ASW, and GC (higher is better for all). Detailed explanations of each metric are provided in Experiment Settings (Section 3). Table 4 offers an aggregated performance summary across three datasets.

**Table 1.** Model performance comparison on the PBMC dataset. Scores indicating the best performance are marked as bold while the second best are marked with underline.

| Model | Alignment Accuracy |  | Embedding Accuracy |  |  |
| --- | --- | --- | --- | --- | --- |
|  | FOSCTTM | CTM | AMI | batch-ASW | GC |
| COOT | 0.109±0.000 | 0.573±0.000 | *NA* | *NA* | *NA* |
| Prior-informed COOT | **0.038**±0.000 | 0.772±0.000 | *NA* | *NA* | *NA* |
| GLUE | <u>0.039</u>±0.003 | <u>0.856</u>±0.021 | **0.781**±0.008 | 0.869±0.024 | 0.965±0.001 |
| scConfluence | 0.048±0.003 | 0.853±0.012 | 0.768±0.007 | 0.840±0.015 | 0.978±0.011 |
| uniPort | 0.080±0.01 | 0.829±0.012 | 0.763±0.003 | 0.856±0.008 | <u>0.979</u>±0.003 |
| MOSCOT | 0.088±0.003 | 0.783±0.003 | 0.739±0.005 | **0.904**±0.003 | 0.941±0.005 |
| LIGER iNMF | 0.080±0.015 | 0.781±0.025 | 0.711±0.020 | 0.836±0.011 | 0.906±0.013 |
| scCotag | 0.040±0.001 | **0.869**±0.004 | <u>0.774</u>±0.006 | <u>0.892</u>±0.005 | **0.980**±0.003 |

**Table 2.** Model performance comparison on the BMMC dataset. Scores indicating the best performance are marked as bold while second best are marked with underline.

| Model | Alignment Accuracy |  | Embedding Accuracy |  |  |
| --- | --- | --- | --- | --- | --- |
|  | FOSCTTM | CTM | AMI | batch-ASW | GC |
| COOT | 0.366±0.000 | 0.234±0.000 | *NA* | *NA* | *NA* |
| Prior-informed COOT | 0.068±0.000 | 0.639±0.000 | *NA* | *NA* | *NA* |
| GLUE | <u>0.049</u>±0.001 | <u>0.731</u>±0.003 | **0.752**±0.004 | **0.937**±0.002 | <u>0.932</u>±0.005 |
| scConfluence | 0.108±0.003 | 0.512±0.012 | 0.609±0.036 | 0.804±0.033 | 0.838±0.034 |
| uniPort | 0.071±0.002 | 0.697±0.006 | 0.689±0.008 | 0.903±0.006 | **0.935**±0.004 |
| MOSCOT | 0.104±0.005 | 0.674±0.006 | 0.663±0.005 | 0.853±0.004 | 0.885±0.006 |
| LIGER iNMF | 0.109±0.013 | 0.629±0.031 | 0.678±0.012 | 0.808±0.025 | 0.848±0.013 |
| scCotag | **0.037**±0.001 | **0.743**±0.004 | <u>0.726</u>±0.06 | <u>0.918</u>±0.004 | 0.919±0.004 |

**Table 3.** Model performance comparison on the Tsai dataset. Scores indicating the best performance are marked as bold while second best are marked with underline.

| Model | Alignment Accuracy |  | Embedding Accuracy |  |  |
| --- | --- | --- | --- | --- | --- |
|  | FOSCTTM | CTM | AMI | batch-ASW | GC |
| COOT | 0.294±0.000 | 0.319±0.000 | *NA* | *NA* | *NA* |
| Prior-informed COOT | <u>0.105±0.000</u> | **0.869±0.000** | *NA* | *NA* | *NA* |
| GLUE | 0.135±0.006 | 0.847±0.014 | 0.753±0.008 | 0.759±0.010 | 0.745±0.009 |
| scConfluence | 0.112±0.003 | 0.786±0.006 | 0.707±0.009 | 0.725±0.015 | 0.712±0.020 |
| uniPort | 0.117±0.004 | 0.835±0.013 | 0.704±0.013 | **0.850±0.006** | <u>0.812±0.025</u> |
| MOSCOT | 0.174±0.003 | 0.718±0.006 | 0.641±0.007 | 0.736±0.011 | 0.687±0.014 |
| LIGER iNMF | 0.175±0.007 | 0.733±0.032 | 0.650±0.017 | <u>0.817±0.008</u> | 0.668±0.038 |
| scCotag | **0.091±0.001** | **0.869±0.001** | **0.761±0.005** | 0.804±0.008 | **0.821±0.004** |

**Table 4.**
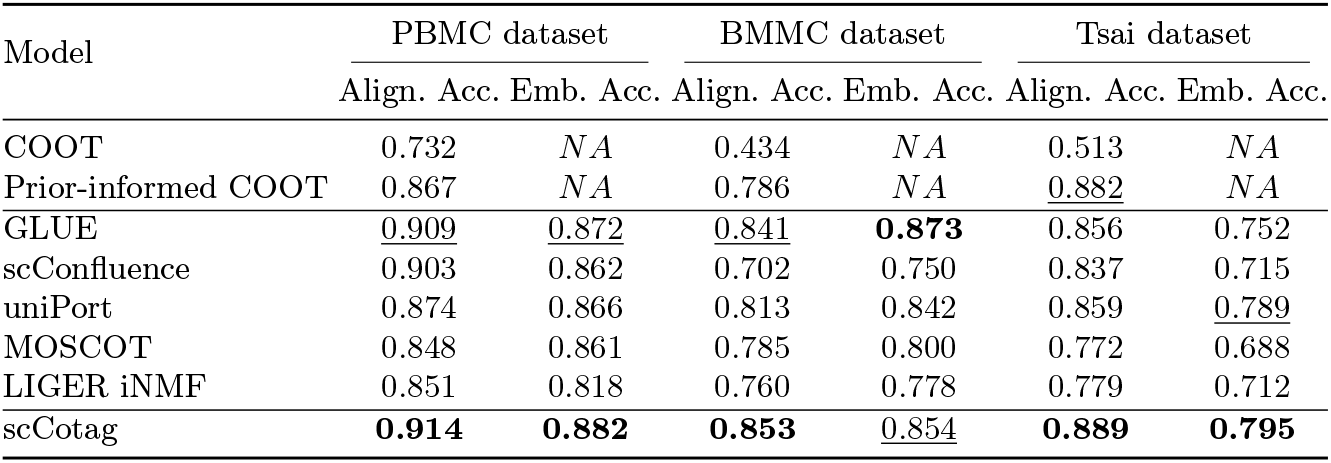
Summarized model performance comparison across three datasets. Embedding accuracy is defined as *Mean(AMI + batch-ASW + GC* ) while alignment accuracy is defined as *Mean((1-FOSCTTM) + CTM)*. Scores indicating the best performance are marked as bold while second best are marked with underline.

**Table 5.**
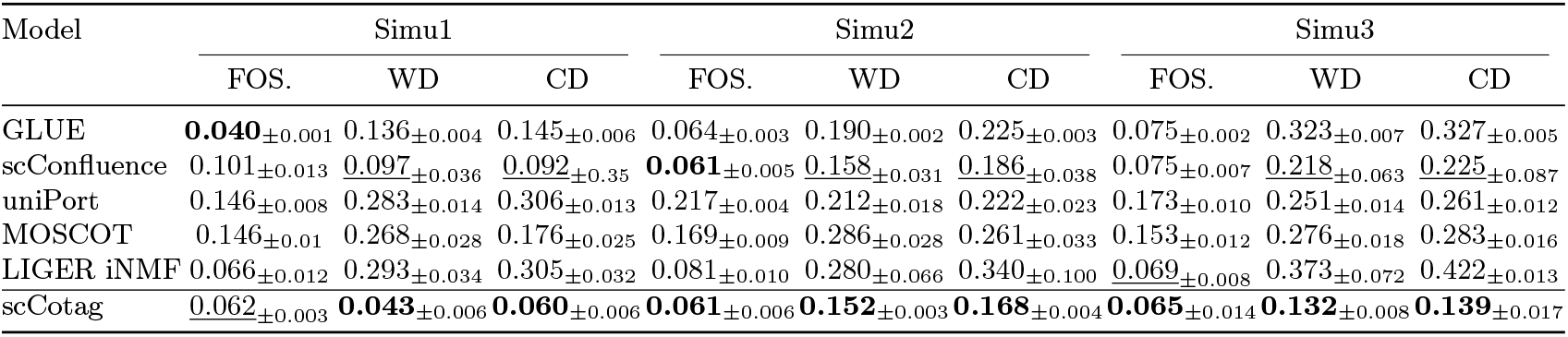
Model performance comparison under three simulated imbalance scenarios. FOS. stands for the FOSCTTM score, WD stands for the within-modality distortion, and CD stands for the cross-modality distortion. Scores indicating the best performance are marked as bold while second best are marked with underline.

## Appendix B: model performance on simulated imbalance datasets

We assessed model performance of scCotag and competing methods under three simulated unbalanced datasets, denoted by *Simi1, Simu2*, and *Simu3* (See Section 3. for dataset curation details). The model performance is evaluating by the alignment accuracy (FOSCTTM) on alignable cells as well as geometry preservation (within-modality distortion and cross-modality distortion, denoted by WD and CD, respectively) on unalignable cells (See Section 3 for metrics definition). For imbalance scenarios, we repeated the experiment five times with different random seed and reported the median and standard deviation as median ± standard deviation. We reported median values here since the model performance exhibited high variability across different seeds.

## Appendix C: Ablation Study

We conducted ablation study to validate that each component in our framework contributes to its superior performance. Specifically, we designed three ablated settings, including *scCotag-NoCellPrior, scCotag-NoFeaturePrior*, and *scCotag-NoPriorCOOT* tested on three paired datasets: PBMC (Table 6), BMMC (Table 7), and Tsai (Table 8). *scCotag-NoCellPrior* is to train the model without utilizing the cell transportation plan for cell representation learning by setting the *λ*_*RegBary*_ = 0 while *scCotag-NoCellPrior* is trained without utilizing the feature transportation plan for graph reweighting. For *scCotag-NoCellPrior*, we replaced the initial cell and feature priors with uniform marginals used in vanilla COOT. In addition, we also compare the model performance between KNN-based geometry preservation and our anchor-based geometry preservation under simulated imbalance scenarios in Table 9.

**Table 6.**
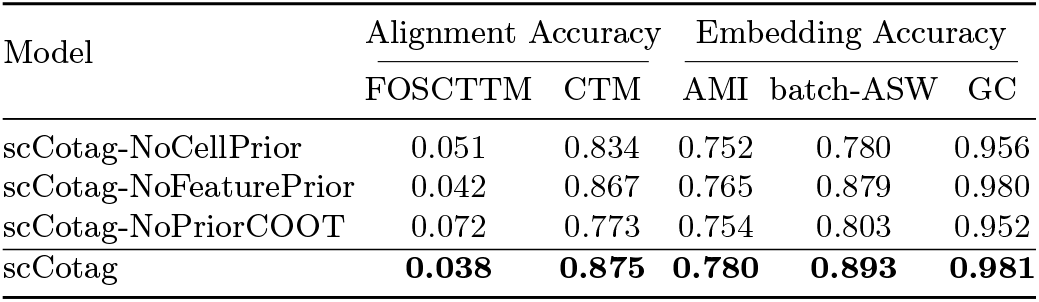
Ablation study on the PBMC dataset. Scores indicating the best model performance are marked as bold.

**Table 7.** Ablation study on the BMMC dataset. Scores indicating the best model performance are marked as bold.

| Model | Alignment Accuracy |  | Embedding Accuracy |  |  |
| --- | --- | --- | --- | --- | --- |
|  | FOSCTTM | CTM | AMI | batch-ASW | GC |
| scCotag-NoCellPrior | 0.052 | 0.695 | 0.698 | 0.847 | 0.911 |
| scCotag-NoFeaturePrior | 0.042 | 0.740 | 0.701 | 0.923 | 0.881 |
| scCotag-NoPriorCOOT | 0.061 | 0.635 | 0.670 | 0.914 | 0.842 |
| scCotag | **0.039** | **0.747** | **0.729** | **0.925** | **0.924** |

**Table 8.** Ablation study on the Tsai dataset. Scores indicating the best model performance are marked as bold.

| Model | Alignment Accuracy |  | Embedding Accuracy |  |  |
| --- | --- | --- | --- | --- | --- |
|  | FOSCTTM | CTM | AMI | batch-ASW | GC |
| scCotag-NoCellPrior | 0.105 | 0.839 | 0.723 | 0.709 | 0.769 |
| scCotag-NoFeaturePrior | 0.097 | 0.848 | 0.737 | **0.814** | 0.789 |
| scCotag-NoPriorCOOT | 0.102 | 0.811 | 0.687 | 0.786 | 0.778 |
| scCotag | **0.091** | **0.869** | **0.760** | <u>0.792</u> | **0.817** |

**Table 9.**
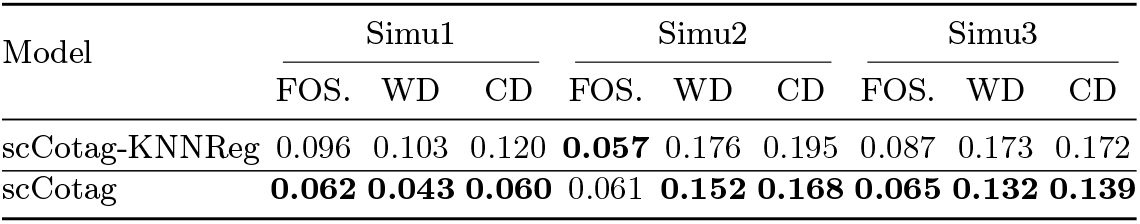
Ablation study under three simulated imbalance scenarios. FOS. stands for the FOSCTTM score, WD stands for the within-modality distortion, and CD stands for the cross-modality distortion. Scores indicating the best model performance are marked as bold.

